# INTERACTION BETWEEN BEES AND CULTIVATED PLANTS: POLLEN CONTRIBUTION IN HONEYS FROM BURKINA FASO

**DOI:** 10.64898/2026.10.15.759302

**Authors:** Oswald Gilbert Dingtoumda, Souhaïbou Sawadogo, Comlan Mawussi Koudegnan, Hugues Roméo Bazie, Antoine Sanon, Zakaria Ilboudo

**Affiliations:** Laboratoire d’Entomologie Fondamentale et Appliquée (LEFA), Unité de Formation et de Recherche en Sciences de la Vie et de la Terre, Université Joseph KI-ZERBO, Ouagadougou 03 BP 7021, Ouagadougou, Burkina Faso; Laboratoire de Recherche Forestière (LRF), Unité de Recherche de la Palynologie, Université de Lomé, 01BP 1515 Lomé 1, Togo; Laboratoire Biosciences (LaBios), Unité de Formation et de Recherche en Sciences de la Vie et de la Terre, Université Joseph KI-ZERBO, Ouagadougou 03 BP 7021, Ouagadougou, Burkina Faso

**Keywords:** : Honeybee, honey, pollen analysis, melliferous plants, phytogeographical zones, Burkina Faso

## Abstract

The domestic honeybee forages on melliferous plants, collecting nectar, pollen, and resin to feed the colony and transform nectar into honey, the principal product of this interaction. This study focused on the palynological characterization of raw honeys from major agricultural zones of Burkina Faso, in order to better identify the floral origin of pollen collected by bees for their needs. Nine (09) raw honey samples from nine (09) apiaries distributed across three (03) phytogeographical zones of Burkina Faso were collected and analyzed. The results revealed 123 pollen taxa grouped into 110 genera and 47 families. Among these identified taxa, six (06) corresponded to cultivated plants: *Abelmochus esculentus* (2.49%), *Arachis hypogaea* (1.17%), *Gossypium hirsitum* (2.92%), *Hibiscus sabdariffa* (2.56%), *Sesamum indicum* (3.97%), and *Vigna unguiculata* (1.51%). The families of Leguminosae and Malvaceae were predominant, representing 22.76% and 8.94% of relative abundance, respectively. Quantitative analysis of pollen grains revealed a contribution ranging from 7.64% to 24.87% per honey sample for pollen originating from cultivated plants. Nevertheless, wild plant species remained the most important pollen sources, with a total relative abundance of 85.35%. Knowledge of the botanical origin of pollen grains collected by bees provides insight into foraging strategies in agricultural environments and allows for the assessment of the effective contribution of each type of resource to colony needs.

## 1. Introduction

Beekeeping is an important sector of the agricultural economy, given the role played by domestic bee populations in pollination and honey production [1]. In Burkina Faso, beekeeping has always held significant socio-economic and environmental importance, due to the income generated by its practice, combined with climatic conditions and the melliferous flora favorable to its development.

It is well established that domestic bees depend almost exclusively on the plant world for their nutrition [2–4]. Indeed, the survival, productivity, and development of any bee colony are closely linked to its foraging activity [5,6]. During this activity, worker bees generally restrict their foraging range to flowers belonging to certain plant species from which they derive their main food resources [7]. The choice of flowers visited depends on the season, floral preference, floral morphology, and the availability of flowers within the foraging range of the hive bees [8,9].

Bees in agricultural environments maintain a close relationship with cultivated plants [10]. Several studies have reported the dependence of domestic bees on cultivated plants in agricultural landscapes during certain periods of the year [11]. Indeed, colonies living in these agroecosystems are highly dependent, both in quantity and quality, on the nectar and pollen resources provided by crop fields [12]. Consequently, the pollen collected and brought back to the hive by bees during the cropping season may originate from either wild or cultivated plant species [13]. According to [14], 87 crops, representing 70% of the 124 crops primarily used in human nutrition worldwide, rely on these pollinating insects.

The plants whose pollen has been foraged by domestic bees can be identified through pollen analysis of honey samples, also known as melissopalynology [15,16]. Melissopalynology is a comprehensive method that, through a single type of analysis namely the examination of pollen extracted from honey provides an overall understanding of bee plant interactions during a given period. This approach is based on the fact that pollen grains contained in natural honeys have been transported by bees and originate from the flowers they visited. The identification of these pollen grains therefore makes it possible to determine the floral and geographical origins of honey, since the pollen spectrum of a honey reflects the vegetation surrounding the hive within a radius of several kilometers.

In Burkina Faso, previous studies on the knowledge and identification of melliferous plants have been conducted [17,18], as well as palynogram analyses of honey samples [19,20]. These a fore mentioned works have documented several melliferous species in the regions concerned. However, research specifically assessing the contribution of pollen from cultivated plants in apiaries established within agricultural landscapes has not been concretely undertaken. The overall objective of the present study is to identify the main plant species serving as foraging sources for domestic bees, whose pollen grains are found in honey originating from agricultural landscapes in Burkina Faso. More specifically, it aims (i) to identify the principal plant species visited by bees for pollen collection and (ii) to evaluate, in proportion, the pollen contribution of cultivated species to domestic bees.

## 2. Materials and Methods

### 2.1. Sampling Sites and Period

Honey sampling for this study was conducted between October and November 2023 in nine (09) apiaries distributed across three phytogeographical zones of Burkina Faso. These included apiaries located in Gourcy and Ouahigouya, belonging to the sub-Sahelian phytogeographical zone; apiaries in Arbollé, Dédougou, and Gassan, belonging to the northern Sudanian phytogeographical zone; and apiaries in Boromo, Houndé, Bobo-Dioulasso, and Orodara, belonging to the southern Sudanian phytogeographical zone (Figure 1).

**Figure 1:**
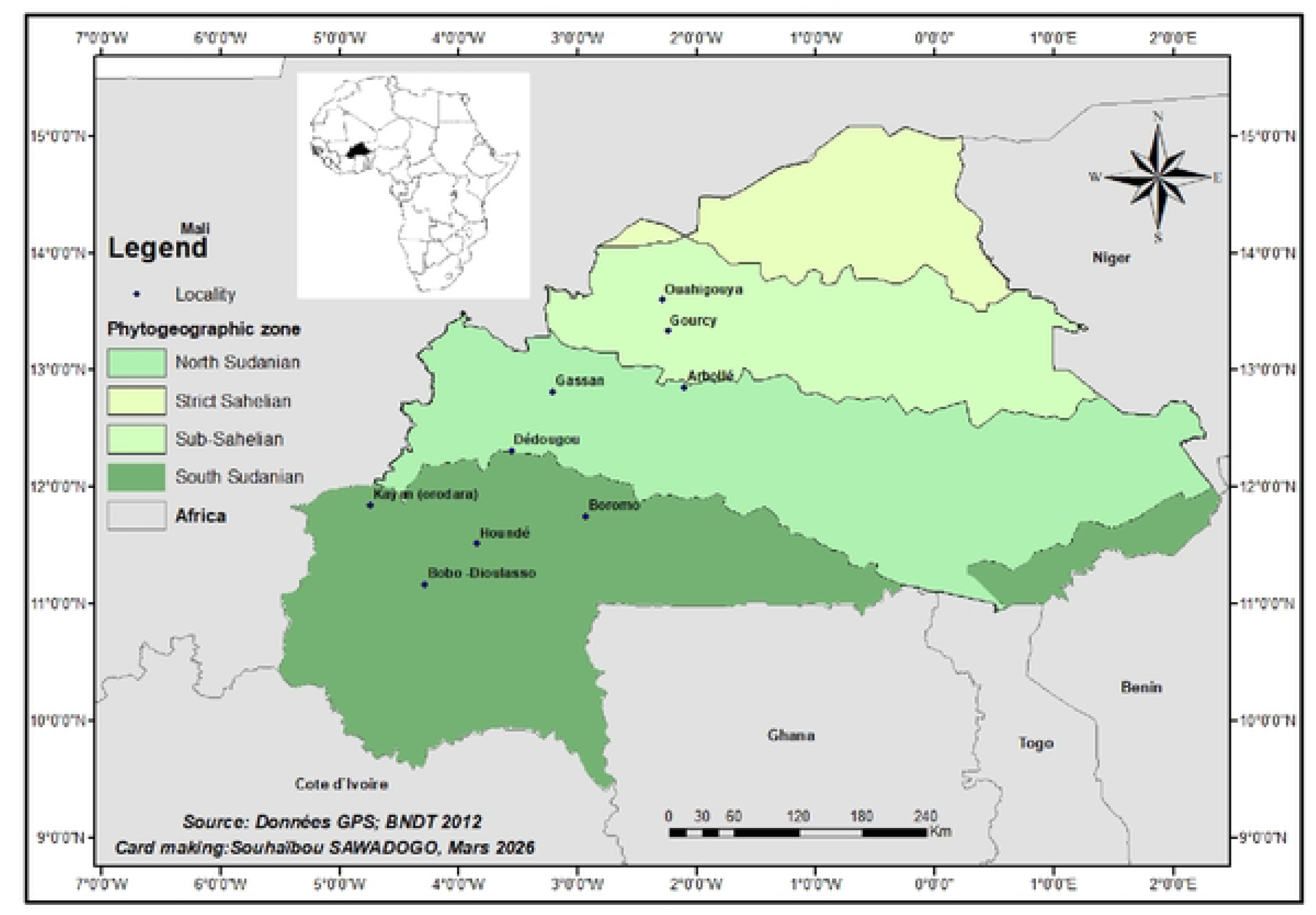
Study area and honey sampling localities The results show that 85.35% of the identified pollen grains originated from wild plants, while 14.65% were derived from cultivated plants (Figure 2).

### 2.2. Biological material

The biological material used for this study consisted of raw honey collected from nine (09) apiaries located in the aforementioned localities. In each apiary, one (01) hive was randomly selected and visited, and a honey sample was taken. Sampling was carried out on combs newly formed by the bees in order to obtain a better assessment of the floral phenology of the foraged plant species. The collected honey samples were coded and stored in sterile, hermetically sealed plastic vials before being transported to the laboratory for further analyses.

### 2.3. Palynological analysis of honey

The palynological analysis of honey samples was carried out in collaboration with the Palynology Research Unit of the Forest Research Laboratory at the University of Lomé, Togo. In the laboratory, the analyses were performed using Erdtman’s acetolysis method [21], revised by [11]. For this purpose, 20 ml of raw honey were collected per sample and processed through successive washings with an acidified solution. This technique aims to remove residues such as proteins, sugars, waxes, and other substances from the pollen membrane, thereby improving the visibility of the exine and enabling the identification of pollen grains from the plant species present in each analyzed honey sample. A volume of twenty (20) μl of the pellet from each sample, obtained after a series of centrifugations, was mounted between a slide and cover slip for the identification and counting of pollen grains using a photonic microscope (Olympus® SC35) at magnifications of 200x and 400x. During microscopic observations, a complete scanning of the entire preparation surface was performed to ensure the detection of the majority of pollen grains present. The identification of pollen grains was carried out to the taxonomic level of species and, where necessary, to the genus or family level. Determinations were made using reference collections and pollen illustration works [22–25] available at the Palynology Research Unit of the Forest Research Laboratory.

### 2.4. Data Processing and Analysis

The classification of the identified taxa was carried out according to the updated phylogenetic system APG IV [26]. Data entry was performed using Microsoft Excel 2016. Graphical representations were generated with the R Studio interface of the R software, version 4.3.1. The relative abundance of the different families and pollen taxa was calculated using the following formulas:

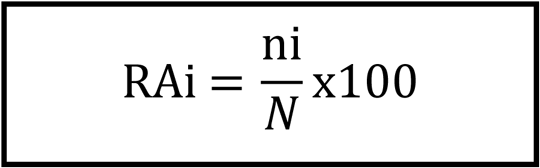

- RAi = relative abundance of taxon *i* (%)
- ni = number of pollen grains belonging to taxon *i*
- N = total number of pollen grains counted in the sample

## 3. Results

### 3.1. Specific richness of the analyzed samples

The specific richness of the honey samples obtained ranged from 28 species (honey from Ouahigouya, OHG) to 48 species (honey from Houndé, HDE). In contrast, pollen richness varied between 538 and 1719 pollen grains. The highest abundance of identified taxa was observed in the sites of Houndé (HDE, 1719 pollen grains), Boromo (BRM, 1513 pollen grains), and Gassan (GSN, 1295 pollen grains). A total of 27 families were predominantly represented in the apiary of Orodara (ODR), compared to only 19 families recorded in the apiary of Ouahigouya (OHG) (Table 1).

**Table 1:** Abundance of plant families and species identified by pollen analysis of honeys from different localities.

| Phytogeographical zone | Localities | Number of families | Number of taxa | Pollen |  |
| --- | --- | --- | --- | --- | --- |
|  |  |  |  | Number | Percentage (%) |
| Sub-Sahelian | GRC | 25 | 41 | 746 | 7.48 |
|  | OHG | 19 | 28 | 808 | 8.10 |
| North Sudanian | ARB | 21 | 38 | 538 | 5.4 |
|  | DDG | 21 | 37 | 1144 | 11.48 |
|  | GSN | 26 | 43 | 1295 | 12.98 |
| South Sudanian | BRM | 23 | 40 | 1513 | 15.17 |
|  | BOB | 21 | 41 | 1033 | 10.35 |
|  | HDE | 23 | 48 | 1719 | 17.24 |
|  | ODR | 27 | 45 | 1176 | 11.8 |
| **Totals** | **9** | **47** | **123** | **9972** | **100** |

*ARB* = *Arbollé*; *GRC* = *Gourcy*; *OHG* = *Ouahigouya*; *BRM* = *Boromo*; *DDG* = *Dédougou*; *GSN* = *Gassan*; *HDE* = *Houndé*; *BOB* = *Bobo*; *ODR* = *Orodara*.

### 3.2. Distribution of species according to families

Table 2 below presents the list of the 47 families to which the identified pollen taxa belong, the number of species per family, as well as the proportions of species obtained by family. The most represented families are Leguminosae (Fabaceae) and Malvaceae, with 28 and 11 plant species identified, respectively (22.76% and 8.94% of the total). They are followed by Meliaceae, Phyllanthaceae, and Rubiaceae, each with 6 species, corresponding to 4.87% of relative abundance. Among the 47 inventoried families, 31 are represented by a single species, each accounting for 0.81% of relative abundance.

**Table 2:** Taxonomic or specific representativeness of the identified families.

| Nº | Families | Number of taxa | Relative abundance (%) | Nº | Families | Number of taxa | Relative abundance (%) |
| --- | --- | --- | --- | --- | --- | --- | --- |
| 1 | Acanthaceae | 1 | 0.81 | 25 | Loranthaceae | 1 | 0.81 |
| 2 | Amaranthaceae | 1 | 0.81 | 26 | Malvaceae | 11 | 8.94 |
| 3 | Anacardiaceae | 5 | 4.07 | 27 | Meliaceae | 6 | 4.87 |
| 4 | Annonaceae | 1 | 0.81 | 28 | Moraceae | 1 | 0.81 |
| 5 | Arecaceae | 4 | 3.25 | 29 | Moringaceae | 1 | 0.81 |
| 6 | Asteraceae | 3 | 2.43 | 30 | Myrtaceae | 3 | 2.43 |
| 7 | Cannabaceae | 1 | 0.81 | 31 | Nymphaeaceae | 1 | 0.81 |
| 8 | Caricaceae | 1 | 0.81 | 32 | Ochnaceae | 1 | 0.81 |
| 9 | Chrysobalanaceae | 1 | 0.81 | 33 | Olacaceae | 1 | 0.81 |
| 10 | Cleomaceae | 1 | 0.81 | 34 | Passifloraceae | 1 | 0.81 |
| 11 | Cochlospermaceae | 1 | 0.81 | 35 | Pedaliaceae | 1 | 0.81 |
| 12 | Colchicaceae | 1 | 0.81 | 36 | Phyllanthaceae | 6 | 4.87 |
| 13 | Combretaceae | 5 | 4.07 | 37 | Proteaceae | 1 | 0.81 |
| 14 | Convolvulaceae | 1 | 0.81 | 38 | Poaceae | 1 | 0.81 |
| 15 | Cordiaceae | 1 | 0.81 | 39 | Polygalaceae | 1 | 0.81 |
| 16 | Cucurbitaceae | 3 | 2.43 | 40 | Portulacaceae | 1 | 0.81 |
| 17 | Cyperaceae | 1 | 0.81 | 41 | Putranjivaceae | 1 | 0.81 |
| 18 | Dilleniaceae | 1 | 0.81 | 42 | Sapotaceae | 1 | 0.81 |
| 19 | Ebenaceae | 1 | 0.81 | 43 | Rhamnaceae | 2 | 1.62 |
| 20 | Euphorbiaceae | 5 | 4.07 | 44 | Rubiaceae | 6 | 4.87 |
| 21 | Heliotropiaceae | 1 | 0.81 | 45 | Rutaceae | 2 | 1.62 |
| 22 | Lamiaceae | 2 | 1.62 | 46 | Vitaceae | 1 | 0.81 |
| 23 | Leguminosae | 28 | 22.76 | 47 | Zygophyllaceae | 1 | 0.81 |
| 24 | Loganiaceae | 1 | 0.81 | **Total** | **47** | **123** | **100** |

### 3.3. Pollen spectra of raw honeys analyzed

The palynological analysis of raw honey samples in this study allowed the identification of 123 pollen taxa distributed across 110 genera and grouped into 47 families (Table 3). Among these pollen taxa, 108 were identified at the species level (87.80%), 10 at the genus level (8.13%), and 5 at the family level (4.07%). Of the 123 identified pollen taxa, six originated from cultivated plants. These included pollen from *A. esculentus*, *A. hypogaea*, *G. hirsitum*, *H. sabdariffa*, *S. indicum*, and *V. unguiculata*.

**Table 3.**
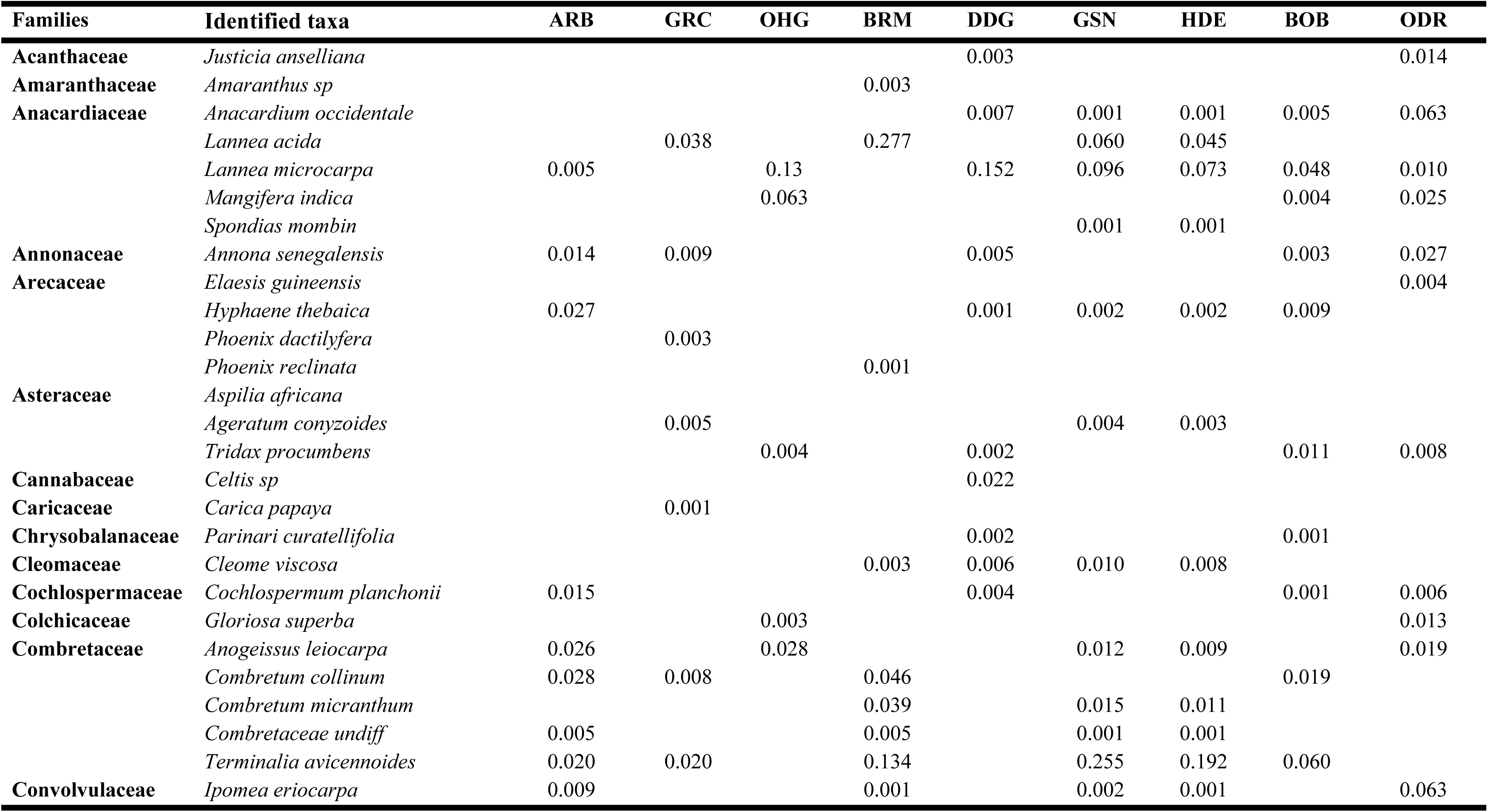
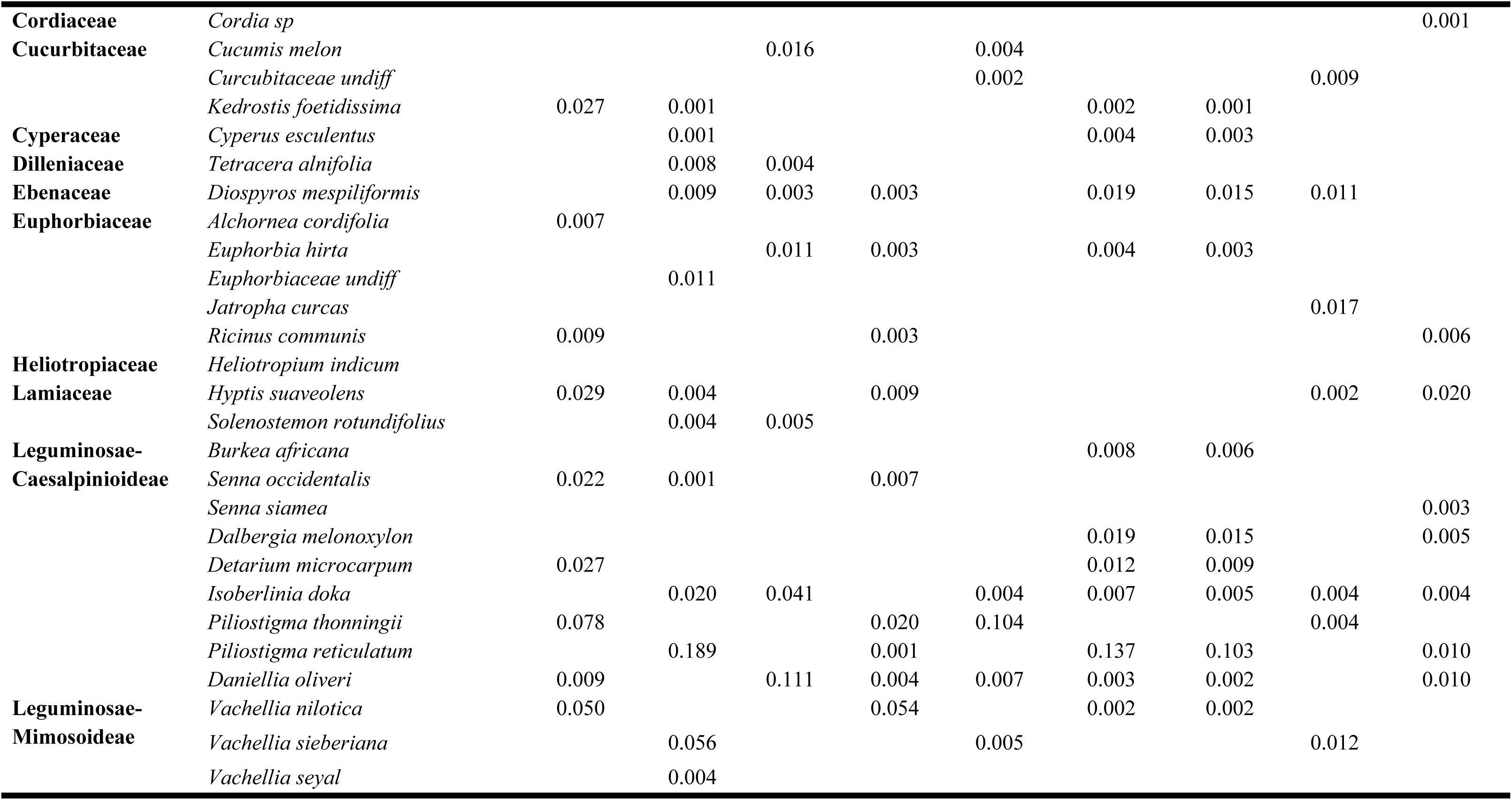
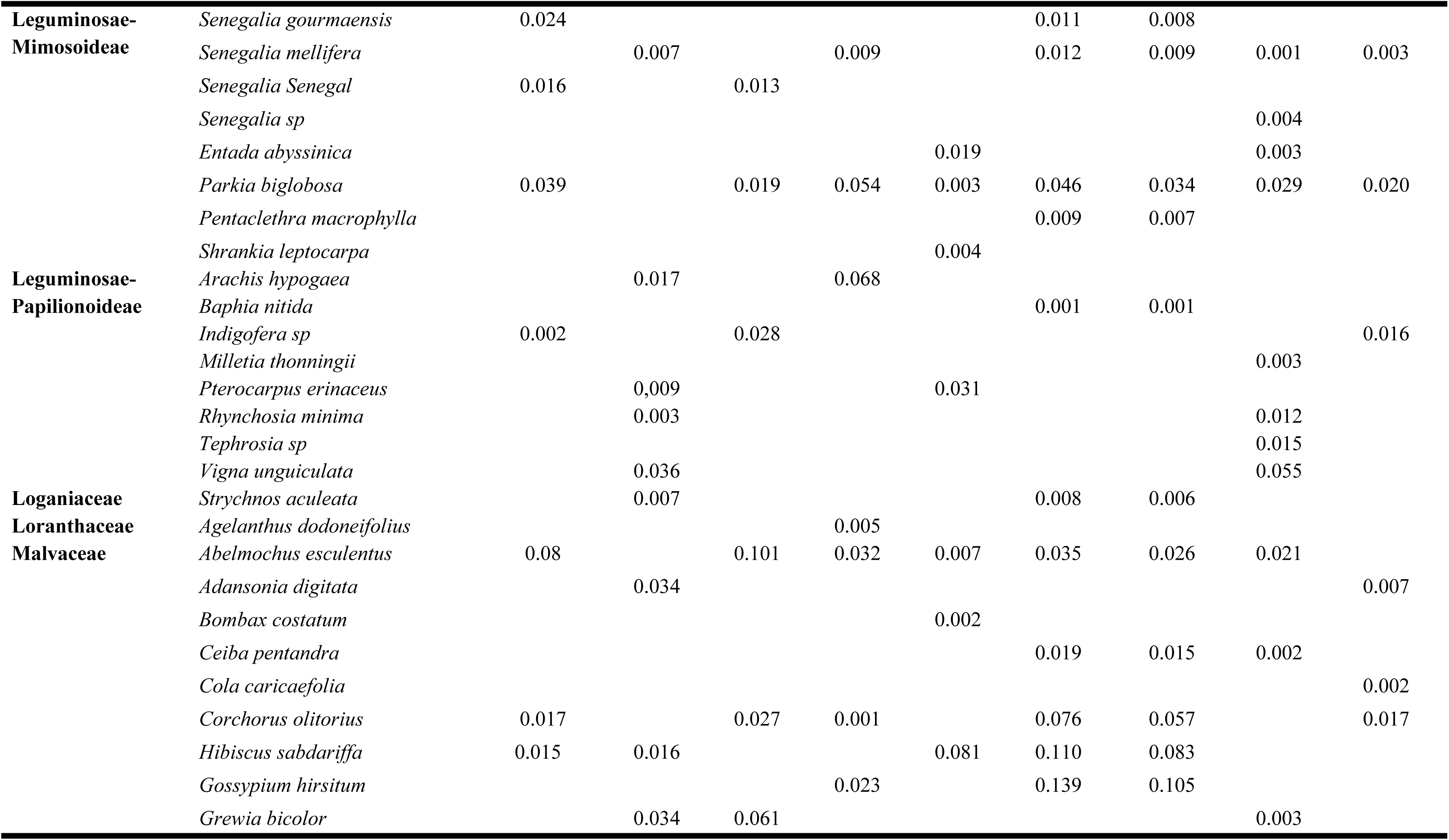
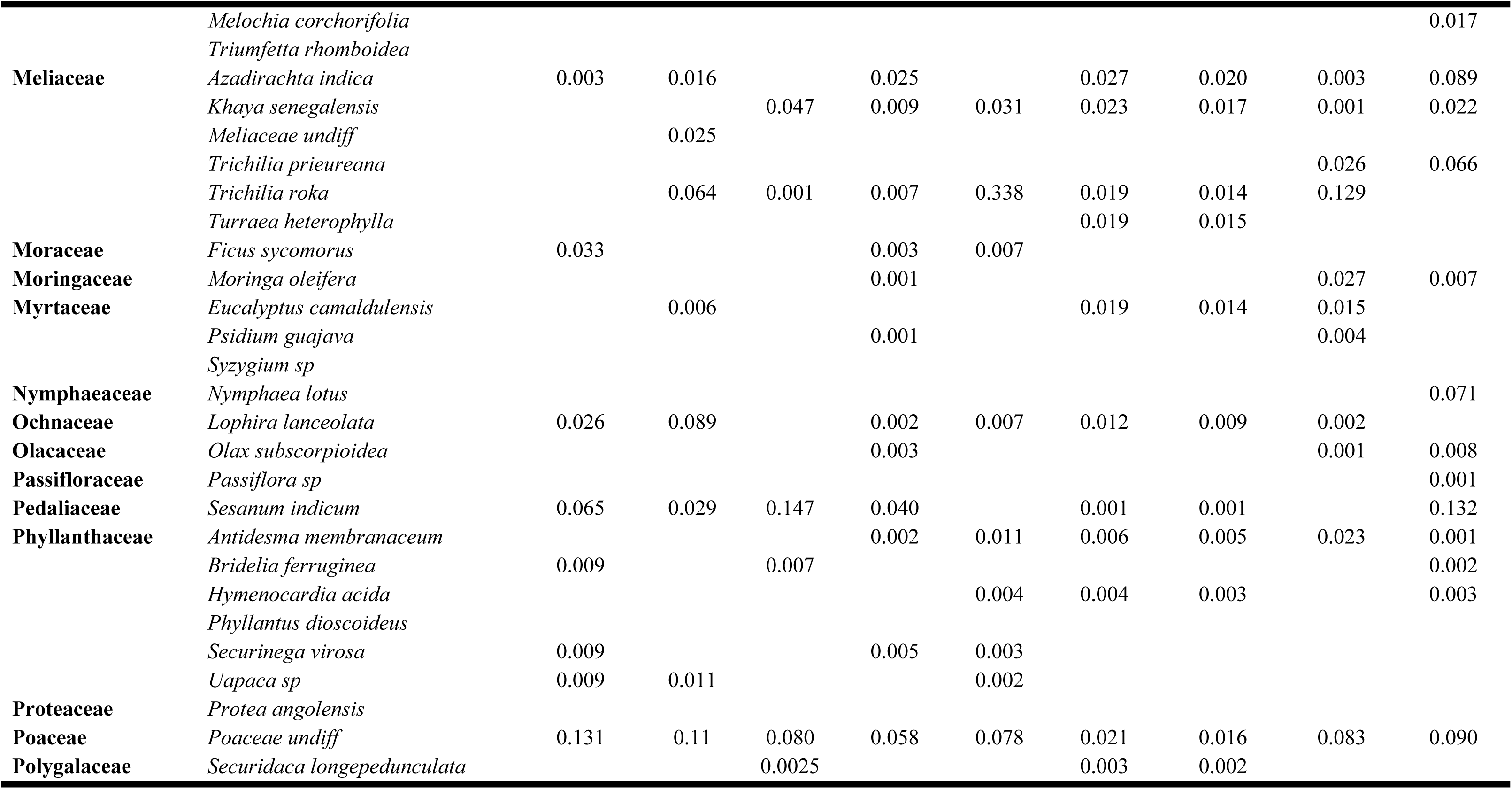
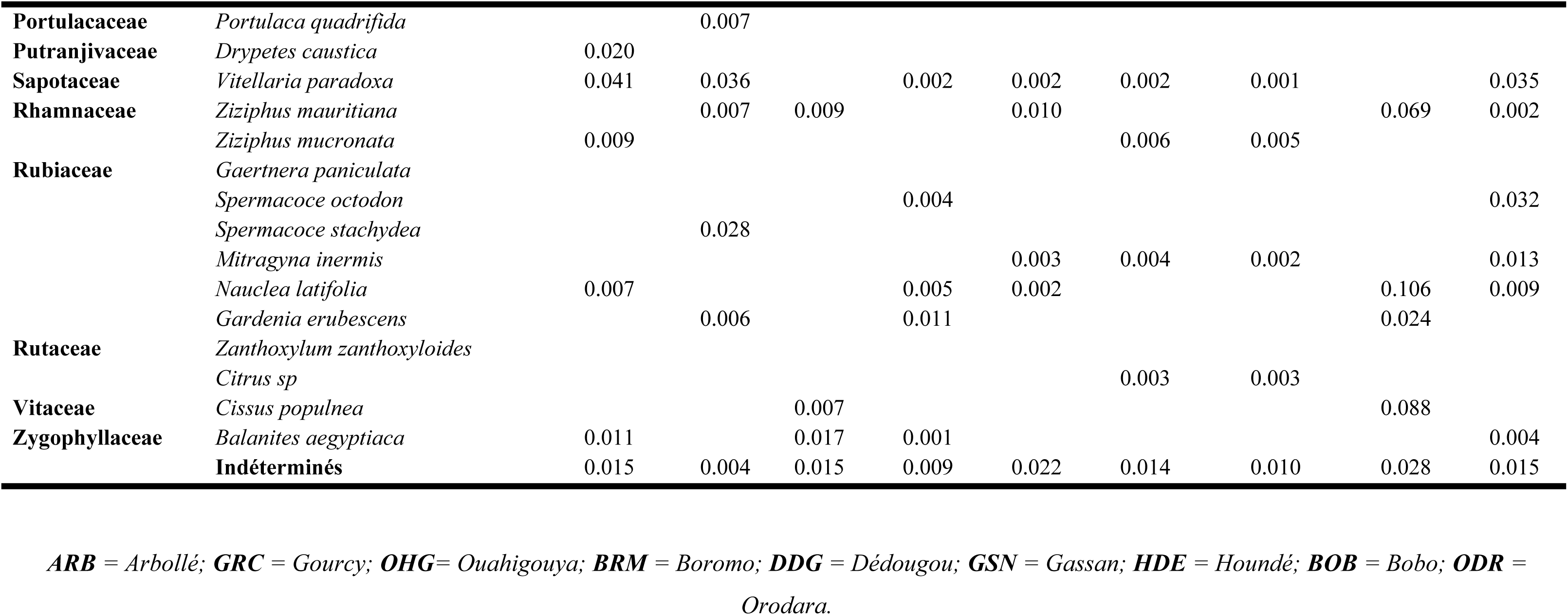
Pollen spectrum of honey samples.

### 3.4. Proportion of pollen taxa according to their floristic origin

The results show that 85.35% of the identified pollen grains originated from wild plants, while 14.65% were derived from cultivated plants (Figure 2).

**Figure 2:**
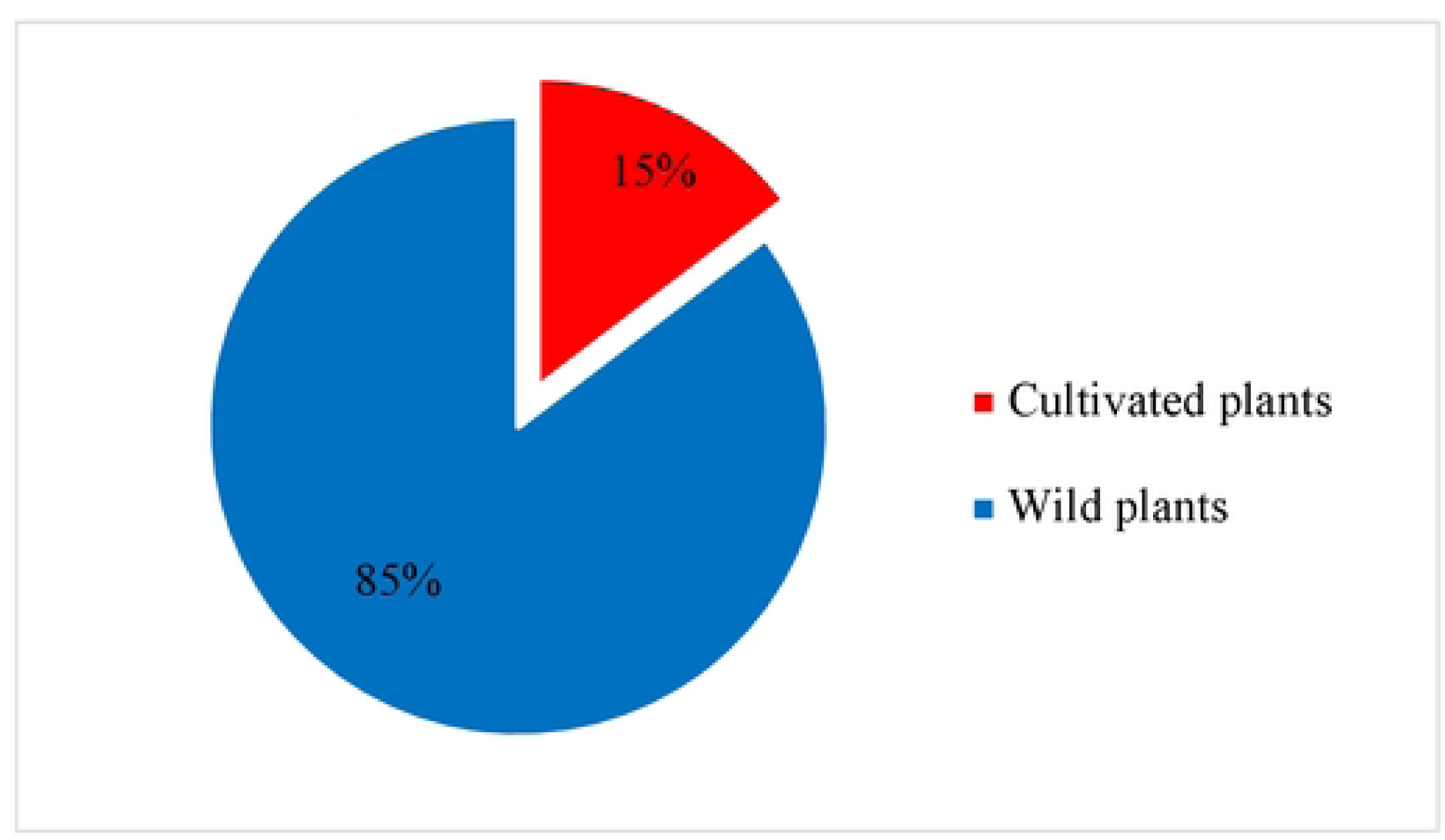
Proportions of taxa according to their origins.

Figure 3 shows images of several pollen grains identified in the raw honey samples.

**Figure 3:**
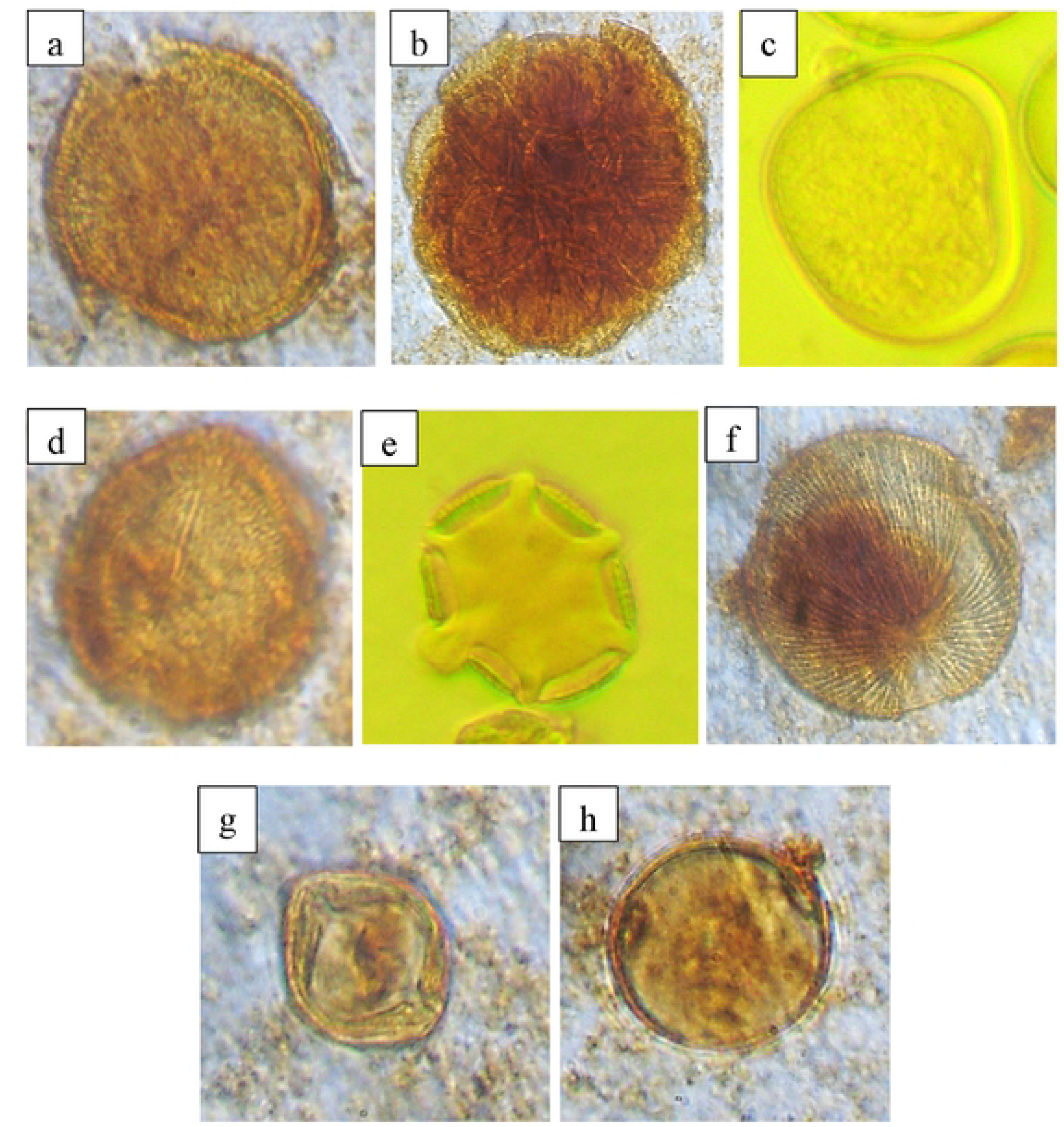
Selected pollen grains from plant species identified in raw honey samples, *Mangifera indica* (a), *Parkia biglobosa* (b), *Poaceae undiff* (c), *Anacardium occidentals* (d), *Hyptis suaveolens* (e), *Isoberlinia doka* (f), *Ricinus communis* (g), *Hymenocardia acida* (h).

### 3.5. Raw abundance of pollen from cultivated plants according to their apicultural origin

The results show that, depending on the apiaries where honey samples were collected, the abundances of pollen from cultivated plants varied in proportion. The highest rate (24.87%) was recorded in the apiary of Ouahigouya. This was followed by the localities of Houndé (21.52%) and Boromo (16.45%). The lowest rate, 7.64%, was obtained in the apiary of Bobo-Dioulasso (Figure 4).

**Figure 4:**
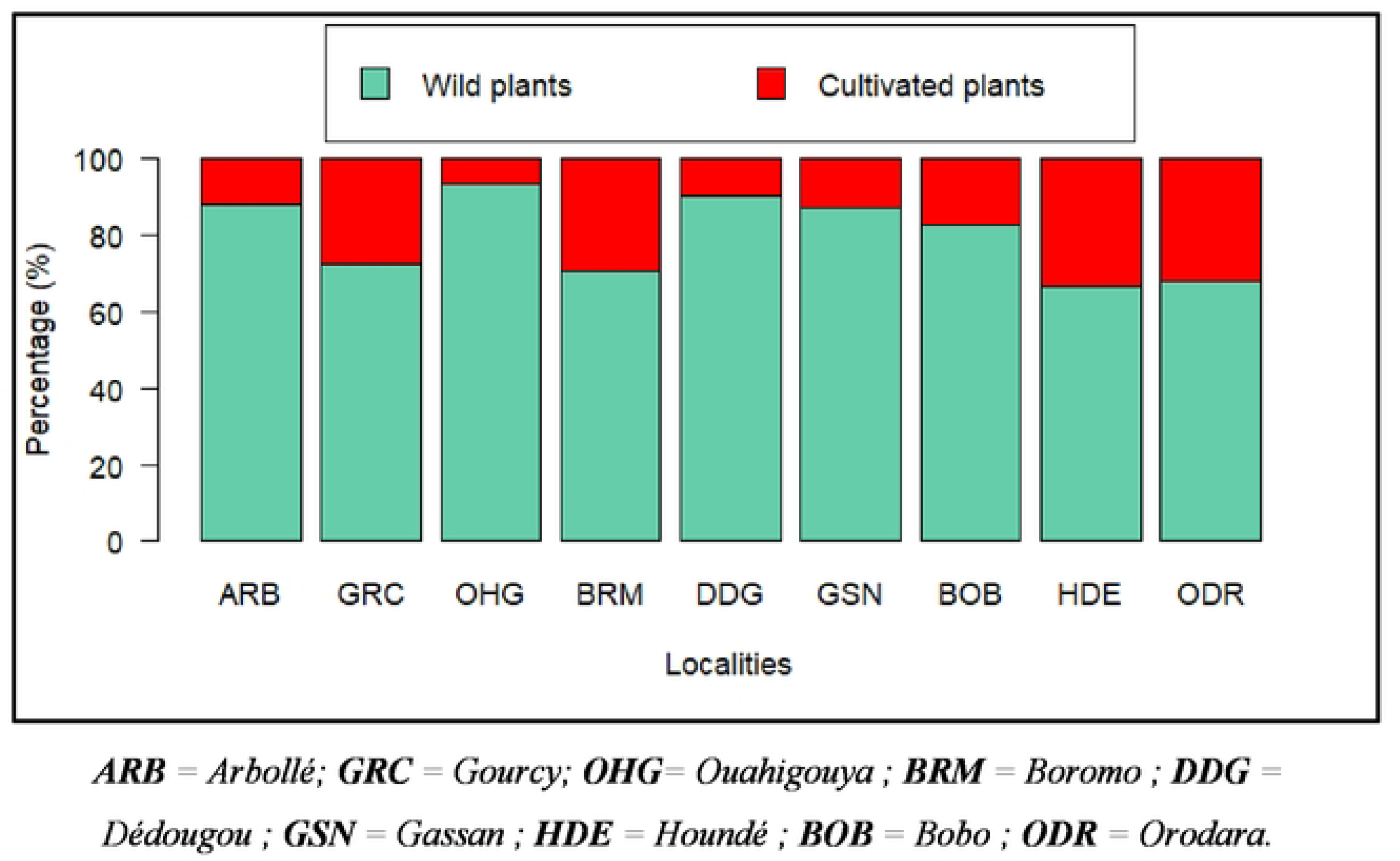
Raw abundance of pollen from cultivated plants across different localities

Figure 5 shows the distribution of pollen taxa from cultivated plants according to the different apiaries of collection. These include pollen from *A. esculentus*, *A. hypogaea*, *G. hirsutum*, *H. sabdariffa*, *S. indicum*, and *V. unguiculata*. In total, pollen from four (04) cultivated plant species was predominantly represented in raw honey samples from the localities of Gourcy (GRC), Boromo (BRM), and Houndé (HDE). The cultivated plant species *Sesamum indicum* was the most widely represented in terms of spatial distribution, occurring in seven (07) out of the nine (09) apiaries sampled. It was followed by *A. esculentus* and *H. sabdariffa*, represented in 6 and 4 sites, respectively.

**Figure 5:**
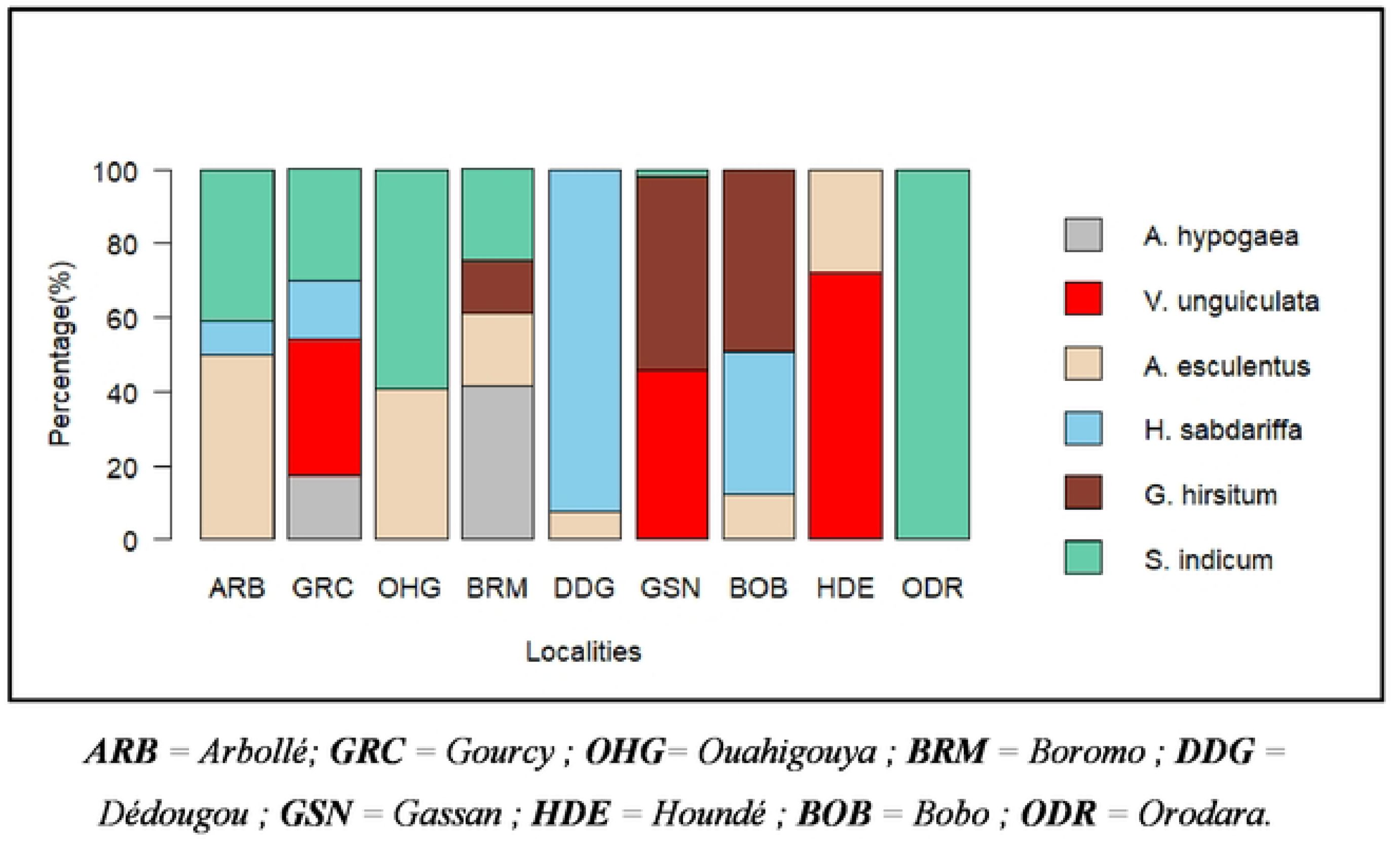
Distribution of pollen taxa from cultivated plants across different localities

## 4. Discussion

Palynological analysis identified 123 pollen taxa distributed across 110 genera and 47 families. The predominant families were Leguminosae (22.76%) and Malvaceae (8.94%). They were the most represented among melliferous taxa, with 28 and 11 taxa respectively. The predominance of species belonging to these families can be explained, on the one hand, by their high abundance in the Sudanian and Sahelian savannas [27,28], and on the other hand by the fact that they include both cultivated and wild species [29]. Furthermore, several studies highlight that these families are particularly rich in nectar and pollen, which confers them a high melliferous value. The predominance of melliferous species belonging to the Leguminosae family has already been reported in Burkina Faso [19,30] as well as in the West African sub-region [11,31,32].

In general, palynological profiles varied from one apiary to another. The number of pollen taxa identified ranged from 28 to 48, obtained respectively in the localities of Ouahigouya and Houndé. The geographical distance between the sampled apiaries could fundamentally explain this variability. Indeed, the apiaries are relatively distant and distributed across the three agro-climatic zones of Burkina Faso. It is well known that melliferous plants vary according to ecological, biotic, and climatic factors [33,34]. This more clearly indicates that the floristic diversity of vegetation formations differs from one locality to another, thereby offering unique floral resources in both quantity and quality. Other factors may also account for the variability observed in palynological profiles. The attractiveness of flowers to honeybees varies according to plant species. It is influenced by floral morphology, phenology, the quality and quantity of pollen, as well as intrinsic factors related to the length of the foragers’ tongue. Flower color, shape, fragrance, and the availability of nectar and/or pollen are all traits that contribute to this attractiveness[35–37]. Finally, the variability in the number of taxa within palynological profiles may also be explained by the presence of numerous minor taxa in the samples. Indeed, 32 out of 47 families are each represented by a single taxon. This confers a unique palynological profile to each honey sample.

The pollen spectrum of the honeys studied confirms the existence of a strong relationship between foraging bees and cultivated plants, accounting for 14.65% of the total abundance. Pollen from six cultivated species was identified in the honey samples from the apiaries studied. These included pollen from *A. esculentus*, *A. hypogaea*, *G. hirsitum*, *H. sabdariffa*, *S. indicum*, and *V. unguiculata*. During the rainy season, the flowering of wild plants remains limited, and pollen and nectar from cultivated plants thus become an essential source for bees, which rely on them to meet the nutritional needs of the colonies [38]. These results confirm the importance of cultivated plants as a temporary foraging source for domestic bees [31,39]. The aforementioned authors demonstrated that cultivated plants account for 64% and 58% of melliferous plant species foraged by bees during the rainy season, respectively. The presence of pollen from these cultivated plants has already been reported in the West African region [12,31].

The proportion of pollen originating from cultivated plants varied between 7.64% and 24.87% depending on the localities. [12] reported values ranging from 0% to 19% in Burkina Faso. The observed differences could be explained by the sampling period and the location of the apiaries in relation to crop fields. Despite the importance of cultivated plants, pollen from wild species remained dominant, accounting for 85.35% of the total relative abundance. This dominance may be attributed to the presence of melliferous taxa whose flowering extends over a long period [13].

Palynological analysis did not allow the identification of pollen grains from the Poaceae family at the species level. However, these pollens accounted for 7.09% of the total relative abundance. Previous studies have shown that certain cultivated Poaceae, such as *Pennisetum glaucum*, *Sorghum bicolor*, and *Zea mays*, may serve as temporary foraging sources for bees, although their melliferous value is relatively low [13]. The bulk of pollen collected by bees generally originates from typically entomophilous plants, but some anemophilous plants, such as most Poaceae, can constitute a temporary source [40]. During the rainy season, many entomophilous plants do not flower, and cultivated Poaceae may therefore emerge as an alternative to compensate for the shortage of pollen resources.

## 5. Conclusion

This study identified the main plant species serving as forage sources for domestic bees and assessed the pollen contribution from cultivated plants through palynological analysis of raw honey. The analysis conducted on nine (09) honey samples revealed 123 pollen taxa grouped into 110 genera and 47 families. The number of identified pollen taxa ranged from 28 to 48. Among the taxa identified, six (06) originated from cultivated plants. The results highlighted a pollen contribution from cultivated plants in all analyzed samples. The proportion of pollen derived from these plants varied between 7.64% and 24.87% depending on the localities, thus revealing a close relationship between honeybees and agricultural crops.

## Data availability

No data was used for the research described in the article

## Acknowledgments

We would like to express our special thanks to the volunteer beekeepers who allowed us to collect honey samples from their apiaries for palynological analyses. We also wish to thank the anonymous reviewer for the valuable comments that helped improve the manuscript.

## Data availability

The authors confirm that all data necessary for supporting the scientific findings of this paper have been provided.

## References

1. STA. Recensement des Apiculteurs et caractérisation des Exploitations Apicoles du BURKINA FASO. 2019.

2. Nguemo DD, Foko J, Pinta JY, Ngouo LV, Tchoumboue J, Zango P. Inventaire et identification des plantes mellifères de la zone soudano-guinéene d’altitude de l’ouest Cameroun. Tropicultura. 2004;22(3):139–45.

3. Brodschneider R, Crailsheim K. Nutrition and health in honey bees. Apidologie. 2010;41(3):278–94.

4. Di Pasquale G, Alaux C, Le Conte Y, Odoux JF, Pioz M, Vaissière BE, Belzunces LP, Decourtye A. Variations in the availability of pollen resources affect honey bee health. PloS one. 2016;11(9): e0162818.

5. Hendriksma HP, Toth AL, Shafir S. Individual and colony level foraging decisions of bumble bees and honey bees in relation to balancing of nutrient needs. Frontiers in Ecology and Evolution. 2019; 7:177.

6. Minarti S, Nursita IW, Puspita PS, Nurwahyuni E. Analysis of Honey Bee (Apis mellifera) Forage Vegetation and Its Implications for Colony Productivity. In: Proceedings of the 5th International Conference on Environmentally Sustainable Animal Industry (ICESAI 2024) [Internet]. Springer Nature; 2025 [cité 3 mai 2026]. p. 422.

7. Auffray V. Nutrition de l’abeille domestique productrice de miel (Apis mellifera) et de sa colonie : revue de la littérature [PhD Thesis]. 2020.

8. Dechaume-Moncharmontp FX. Butinage collectif chez l’abeille Apis mellifera L. : étude théorique et expérimentale. Butinage collectif chez l’abeille *Apis mellifera* L (2003) [Internet]. 2003 [cité 19 avr 2026].

9. Hernández JC, García JE, Wells H, Amaya-Márquez M. Honey Bee Foraging Decisions Are Shaped by Floral Trait Distinctiveness and Perception of Gains or Losses. Insects. 2025;16(9):884.

10. Stein K, Coulibaly D, Stenchly K, Goetze D, Porembski S, Lindner A, Konaté S, Linsenmair EK. Bee pollination increases yield quantity and quality of cash crops in Burkina Faso, West Africa. Scientific Reports. 2017 ;7(1) :17691.

11. Koudegnan CM, Edorh TM, Guelly AK, Batawilla K, Akpagana K. Inventaire des taxons polliniques des miels de la zone guinéenne du Togo : Cas des zones ecofloristiques IV et V. European Scientific Journal [Internet]. 2012 [cité 4 mai 2026] ;8(26).

12. Cencetti T, Orioli L, Nombré I, Di Marco G, Leonardi D, Gismondi A, Canini A. Explorative socio-environmental survey for honey quality assessment in six target provinces of Burkina Faso. Journal of Agriculture and Environment for International Development (JAEID). 2017 ;111(2):361–82.

13. Piroux M. Ressources pollinifères et mellifères de l’Abeille domestique, Apis Mellifera, en paysage rural du nord-ouest de la France [PhD Thesis] [Internet]. Université Blaise Pascal-Clermont-Ferrand II; 2014 [cité 3 mai 2026].

14. Klein AM, Vaissière BE, Cane JH, Steffan-Dewenter I, Cunningham SA, Kremen C, Tscharntke T. Importance of pollinators in changing landscapes for world crops. Proceedings of the royal society B: biological sciences. 2007 ;274(1608) :303–13.

15. Lobreau-Callen D, Darchen R, Le Thomas A. Apport de la palynologie à la connaissance des relations abeilles/plantes en savanes arborées du Togo et du Bénin. Apidologie. 1986 ;17(4) :279–306.

16. Koudegnan CM. Contribution de la palynologie à la caractérisation des miels et à la connaissance de la flore mellifère de la zone guinéenne du Togo. Th Doc, Univ Lomé, Togo. 2015.

17. Guinko S, Guenda W, Millogo Rasolodimby J, Tamini Z, Zoungrana I. Etude des plantes mellifères dans l’ouest du Burkina Faso (Provinces du Houet, de la Comoé et du Kénédougou). [Internet]. 1987 [cité 12 août 2026].

18. Nombre I. Etude des potentialités mellifères de deux zones du Burkina Faso Garango (Province du Boulgou) et Nazinga (Provinche du Nahouri). 2003.

19. Cencetti T, Lippi MM, Nombré I, Orioli L. Pollen analysis of some Burkina Faso honey samples. Webbia. 3 juill 2019;74(2):373–81. doi:10.1080/00837792.2019.1627455

20. Schweitzer P, Nombré I, Kwame A, Boussim JI. Spectrum of plant species foraged by *Apis mellifera adansonii* Latreille in the North Sudanian phytogeographical region of Burkina Faso. Grana. 2 janv 2014;53(1):62–8. doi:10.1080/00173134.2013.853830

21. Erdtman G. Pollen walls and angiosperm phylo-geny. [Internet]. 1960 [cité 4 mai 2026]. Disponible sur: https://www.cabidigitallibrary.org/doi/full/10.5555/19601603471

22. Sowunmi MA. Pollen Grains of Nigerian Plants: I. Woody Species. Grana. déc 1973;13(3):145–86. doi:10.1080/00173137309429891

23. Sowunmi MA. Pollen of Nigerian Plants: II Woody Species. Grana. avr 1995;34(2):120–41. doi:10.1080/00173139509430002

24. Ybert JP. Atlas de pollens de Côte d’Ivoire [Internet]. IRD Editions ; 1979 [cité 11 mai 2026].

25. Bonnefille R, Riollet G. Pollens des savanes d’Afrique orientale [Internet]. Éditions du Centre national de la recherche scientifique Paris; 1980 [cité 11 mai 2026]. Disponible sur: https://library.wur.nl/WebQuery/titel/549790

26. Angiosperm Phylogeny, Chase MW, Christenhusz MJ, Fay MF, Byng JW, Judd WS, Soltis DE, Mabberley DJ, Sennikov AN, Soltis PS. An update of the Angiosperm Phylogeny Group classification for the orders and families of flowering plants: APG IV. Botanical journal of the Linnean Society. 2016;181(1):1–20.

27. Qasim M, Porembski S, Sattler D, Stein K, Thiombiano A, Lindner A. Vegetation structure and carbon stocks of two protected areas within the South-Sudanian Savannas of Burkina Faso. Environments. 2016;3(4):25.

28. Zoungrana BJB, Dimobé K. NDVI-derived vegetation trends and driving factors in West African Sudanian savanna. Am J Plant Sci. 2023;14(10):1130–45.

29. Bayer C. Malvaceae In: Kubitzki, K. & Bayer, C.(eds) The Families and Genera of Vascular Plants V: Flowering plants. Dicotyledons: Malvales, Capparales and Non-betalain Caryophyllales. *Springer-Verlag*, Berlin ; 2003.

30. Kanazoé IW, Nombré I, Vereecken J. Caractérisation de l’apiculture dans les villages riverains du corridor n 1 du Complexe Pô-Nazinga-Sissili (PONASI) dans le Centre-Sud du Burkina Faso. Geo-Eco-Trop. 2021;45(3):455–66.

31. Iritié BM, Wandan EN, Paraiso AA, Fantodji A, Gbomene LL. Identification des plantes mellifères de la zone agroforestière de l’Ecole Supérieure Agronomique de Yamoussoukro (Côte d’Ivoire). European scientific journal. 2014 ;10 (30).

32. Ahouandjinou ST, Yedomonhan H, Adomou AC, Tossou MG, Akoegninou A. Caractéristiques techniques et importance socio-économique de l’apiculture au Nord-Ouest du Bénin: cas de la commune de Cobly. International Journal of Biological and Chemical Sciences. 2016;10(3):1350–69.

33. Aragón-Gastélum JL, Ramírez-Albores JE, Pérez-Suárez M, Vargas-Contreras JA, Aguirre-Crespo FJ, Plascencia-Escalante FO, Serrano-Rodríguez A, Plasencia-Vázquez AH. Assessment of the Impact of Climate Change on the Potential Distributions of Melliferous Plant Species on the Yucatan Peninsula, Mexico: Implications for Conservation Planning. Conservation. 2025;5(3):44.

34. Kanazoé IW, Nombré I, Zoungrana A, Boussim JI. Land-Use Effects on Melliferous Woody Flora in Sudanian Agroforestry Parkland and Protected Area of West Africa. Ecology and Evolution. avr 2026;16(4):e73555. doi: 10.1002/ece3.73555

35. Baude M, Muratet A, Fontaine C, Pellaton M. Plantes et pollinisateurs observés dans les terrains vagues de Seine-Saint-Denis. Observatoire Départemental de la Biodiversité Urbaine (COBU), 65p. 2011.

36. Muth F, Francis JS, Leonard AS. Bees use the taste of pollen to determine which flowers to visit. Biology Letters [Internet]. 2016 [cité 4 mai 2026];12(7).

37. Rowe L, Gibson D, Bahlai CA, Gibbs J, Landis DA, Isaacs R. Flower traits associated with the visitation patterns of bees. Oecologia. juin 2020;193(2):511–22. doi:10.1007/s00442-020-04674-0

38. Letsyo E, Ameka G. Major plants foraged by bees for honey production in Ghana: mapping of bee floral sources for the development of the apicultural industry. Grana. 2 nov 2019;58(6):472–82. doi:10.1080/00173134.2019.1629622

39. Pinta JY, Choumboue J, Dongock N, Zango P, Sitcheu G. Some characteristics of melliferous plants of the western High Lands of Cameroon. In: Proceedings of the 9th Annual Conference of Bioscience, Yaoundé, Cameroon. 2001.

40. Lokossou SC, Tchobo FP, Djossou AJ, Yedomonhan H, Akoegninou A. Diversité et vertus thérapeutiques des ressources floristiques de production de miel au Bénin. Int J Biol Chem Sci. 2019;13(1):294–310.

